# PINK1 regulated mitophagy is evident in skeletal muscles

**DOI:** 10.1101/2023.11.09.566402

**Authors:** Francois Singh, Lea Wilhelm, Alan R. Prescott, Kevin Ostacolo, Jin-Feng Zhao, Margret H. Ogmundsdottir, Ian G. Ganley

**Affiliations:** MRC Protein Phosphorylation and Ubiquitylation Unit, University of Dundee, Dundee DD1 5EH, UK; Department of Physiology, Biomedical Center, School of Health Sciences, University of Iceland, Vatnsmýrarvegur 16, 101 Reykjavik, Iceland; Dundee Imaging Facility, School of Life Sciences, University of Dundee, Dundee DD1 5EH, UK; Department of Physiology, Biomedical Center, Faculty of Medicine, University of Iceland, Sturlugata 8, 101, Reykjavik, Iceland

**Keywords:** Parkinson’s, PINK1, mitophagy, Mutator, muscle

## Abstract

PINK1, mutated in familial forms of Parkinson’s disease, initiates mitophagy following mitochondrial depolarization. However, it is difficult to monitor this pathway physiologically in mice as loss of PINK1 does not alter basal mitophagy levels in most tissues. To further characterize this pathway *in vivo*, we used *mito*-QC mice in which loss of PINK1 was combined with the mitochondrial-associated POLG^D257A^ mutation. We focused on skeletal muscle as gene expression data indicates that this tissue has the highest PINK1 levels. We found that loss of PINK1 in oxidative hindlimb muscle significantly reduced mitophagy. Of interest, the presence of the POLG^D257A^ mutation, while having a minor effect in most tissues, restored levels of muscle mitophagy caused by the loss of PINK1. Although our observations highlight that multiple mitophagy pathways operate within a single tissue, we identify skeletal muscle as a tissue of choice for the study of PINK1-dependant mitophagy under basal conditions.

## Introduction

Mutations of PTEN induced kinase 1 (PINK1/PARK6) result in the development of autosomal recessive forms of early-onset Parkinsońs disease (PD) [1]. PD is among the most common neurodegenerative disorders in the world and is characterized by the selective degradation of dopaminergic (DA) neurons within the substantia nigra pars compacta (SNpc). However, the exact mechanisms leading to this event remain elusive. Mounting evidence suggests that dysregulated mitophagy (the autophagy of mitochondria) could play a central role in the etiology of PD [2]. Most of what we know about the regulation of mitophagy comes from the extensive *in vitro* study of the PINK1/Parkin (PARK2) pathway. PINK1 is stabilized and activated upon mitochondrial depolarization and phosphorylates both ubiquitin and Parkin at their Ser65 residue [3,4]. This leads to a feed-forward mechanism of ubiquitylation of outer mitochondrial membrane (OMM) proteins that results in recruitment of the autophagy machinery, engulfment into autophagosomes and concomitant degradation and recycling in lysosomes. Elucidation of this mechanism has necessitated the use of somewhat artificial conditions in cell lines using protein overexpression and harsh mitochondrial toxicants [5,6]. Given this, the physiological relevance of this pathway is still enigmatic [7].

We previously reported, using the *mito*-QC mouse model, that PINK1 and Parkin do not significantly impact basal mitophagy in several tissues, including in PD related neuronal types [8,9]. This result is consistent with observations from other model organisms, including Drosophila [10] and zebrafish [11] and was key in showing that under basal conditions other mitophagy pathways operate in tissues. Given the nature of the PINK1 pathway, as revealed from the *in vitro* studies mentioned earlier, it is likely that it requires distinct, as yet unknown, stressors to become dominant *in vivo*. Given this, it would be advantageous to identify tissues and conditions where complete flux through the PINK1-dependent mitophagy pathway could be monitored, as this would not only shed insights into how PINK1-dependent mitophagy could go awry in PD, but also provide a setting whereby novel therapeutic approaches could be easily tested in a physiological setting to determine engagement of this mitophagy pathway. In this endeavor, we took two approaches based on our previously well characterized *mito*-QC reporter system and mouse model [8,9,12–16]. The *mito*-QC reporter mouse constitutively expresses an mCherry-GFP tag fused to the OMM. Under normal conditions the mitochondrial network fluoresces both red and green but upon mitophagy, where a mitochondrion is delivered to the lysosome, the acidic microenvironment quenches the GFP fluorescence, but not mCherry. Hence, the proportion of mitophagy within a specific tissue or cell type can accurately be determined by measuring the proportion and size of mCherry-only puncta (mitolysosomes). Firstly, we crossed PINK1 knockout (KO) *mito*-QC mice with mutator mice. The mutator mice (POLG^D257A^) are a commonly used model for mitochondrial dysfunction as they display a mutation in the proofreading exonuclease domain of the DNA polymerase γ gene, leading to the accumulation of mitochondrial DNA mutations [17,18]. Consequently, these mice undergo premature aging, have a reduced lifespan and display sarcopenia and cardiomyopathy [19,20]. Importantly, loss of Parkin has been shown to synergize with mitochondrial dysfunction present in mutator mice resulting in increased DA neuron cell death in the SNpc [18]. Secondly, we reasoned that PINK1-dependent mitophagy will most likely be detectible in tissues where PINK1 expression is the highest. Based on analysis of the Genotype-Tissue Expression project (https://www.gtexportal.org) we focused on skeletal muscle. We observed an effect of PINK1 ablation on mitochondrial content and on basal mitophagy levels in skeletal muscles, with a gradient-like effect of reduced mitophagy dependent on the known oxidative status of distinct muscle fiber types. Of interest, while the mutator mice displayed a global loss of oxidative fibers, the presence of the POLG^D257A^ mutation resulted in restoration of mitophagy in PINK1-null muscles to levels comparable with the wild-type control group. Our results provide strong evidence for PINK1 in contributing to skeletal muscle mitophagy and highlight an existing compensatory mechanism(s) in response to additional mitochondrial stress.

## Results

### Mitochondrial homeostasis is disrupted in POLG^D257A^ and PINK1 skeletal muscles

We bred and analyzed 6-month-old mice that were either homozygous for the POLG^D257A^ mutation, PINK1 KO, or a combination of both. All mice were homozygous for the *mito*-QC reporter. As it has been previously noted [18,21], the presence of the POLG^D257A^ mutation resulted in lower body weight (Figure S1A). We also confirmed loss of PINK1 in immunoprecipitates from whole brain lysates (Figure S1B). To identify tissues that express high levels of PINK1, which are more likely to display effects on mitophagy under basal conditions, we analyzed RNAseq data generated by the Genotype-Tissue Expression (GTEx) project. This resource contains human gene expression data in 54 tissue sites from 948 individuals. Searching the database for PINK1, as well as PRKN, TBK1, OPTN and SQSTM1 (all proteins known to operate in this pathway [22]) revealed that these proteins had the highest expression levels overall in skeletal muscle (Figure 1A). We therefore focused our efforts on this tissue. We examined three types of hindlimb skeletal muscles, based on their well characterized mitochondrial content and metabolic phenotype (glycolytic vs. oxidative). We compared phenotypes in the low oxidative/ high glycolytic part of the gastrocnemius muscle, known as the white gastrocnemius (WGC), with the high oxidative/low glycolytic red and mixed gastrocnemius and plantaris, termed red/mixed calf muscles (RCM). We also examined the highly oxidative hindlimb soleus muscle (SOL). A representative image of these muscle groups from *mito*-QC mice is shown in Figure 1B, which clearly demonstrates the oxidative nature of the soleus as evidence by increasing mitochondrial content (visualized indirectly by the GFP channel of the *mito*-QC reporter). Overall, no significant differences in muscle weight were found between all the genotype groups for both the total gastrocnemius and plantaris, and for the soleus (Figure 1C&D). However, when sub-dissecting the gastrocnemius and plantaris into their white and red/mixed regions, we noticed a significant difference in their relative proportions in POLG^D257A^ mutant mice. We measured the respective weights of these muscle regions and observed a higher mass for the WGC in the POLG^D257A^ groups, suggesting a potential loss of oxidative fibers compared to the control group in the whole muscle (Figure 1C compared with E&F). To confirm this apparent expansion of WGC at the expense of RCM, we visualized cross-sections through whole calf muscles and measured the proportional area of the WGC and of the RCM, based on the relative expression of the *mito*-QC reporter (Figure 1G-H). Confirming our macroscopic observations, these data revealed a loss of the proportion of oxidative fibers in the gastrocnemius of POLG^D257A^ mice.

**Figure 1.**
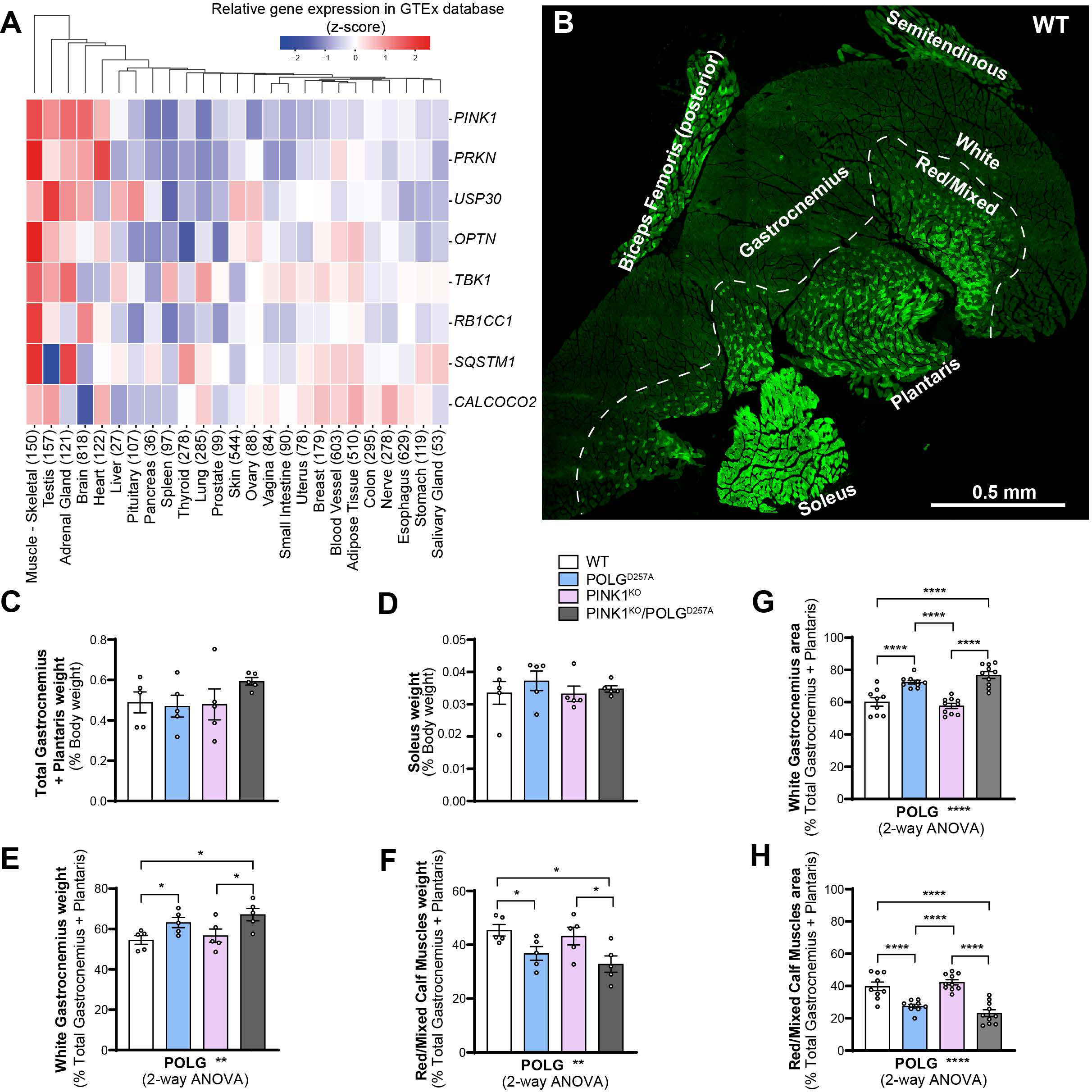
The PINK1 pathway is highly expressed in skeletal muscles and POLG^D257A^ alters calf muscle fibre proportion and mitochondrial content. (A) Gene-expression RNA-seq data analysis of the PINK1/Parkin pathway in different human tissues, obtained from GTEx (Genotype-Tissue Expression). (B) Representative composite tile-scan micrograph of a cross-section from the hindlimb calf skeletal muscles from a *mito*-QC wild-type mouse. Muscles are easily distinguishable by mitochondrial content using the *mito*-QC GFP expression, with the soleus being highly oxidative, the white gastrocnemius being highly glycolytic, and the red/mixed gastrocnemius presenting an intermediate/oxidative phenotype. Dashed lines delimitate the white gastrocnemius from the red/mixed gastrocnemius. Scale bar: 0.5 mm. Relative weights of the (C) total gastrocnemius and plantaris, and (D) soleus expressed proportionally to body weight of each mouse (n=5 per group) of WT, PINK1 knock-out, mutator and double mutant (PINK1^KO^/POLG^D257A^) *mito*-QC mice. Relative weight of the (E) white gastrocnemius (WGC), and (F) red/mixed calf muscles (RCM, red/mixed gastrocnemius + plantaris) expressed proportionally to the weight of the total gastrocnemius + plantaris (n=5 per group). Relative cross-sectional area of the (G) white Gastrocnemius and of the (H) red/mixed calf muscles (red/mixed gastrocnemius + plantaris) (n=9-10). Overall data is represented as mean +/-SEM. Statistical significance is displayed as *p<0.05, and ****p<0.0001.

Regardless of the change of WGC to RCM ratios in POLG^D257A^ mice, we next analyzed mitochondrial content within these areas based on the expression of the *mito*-QC reporter (Figure S1C), and on protein expression (Figure 2). In general, mitochondrial proteins such as HSP60, MNF2 and COXIV were present at a higher level in PINK1 KO tissue, with this increase more pronounced in the oxidative RCM tissues. POLG^D257A^ also caused changes in the mitochondrial proteins, but with the exception of COX IV, the changes were different depending on protein and tissue areas analyzed (Figure 2). This implies a more complex regulation that requires more work to understand. COX IV levels were dramatically reduced in the background of the POLG^D257A^ mutation (regardless of PINK1 status). This phenomenon was more pronounced in WGC and has previously been described in the quadriceps [20,23].

**Figure 2.**
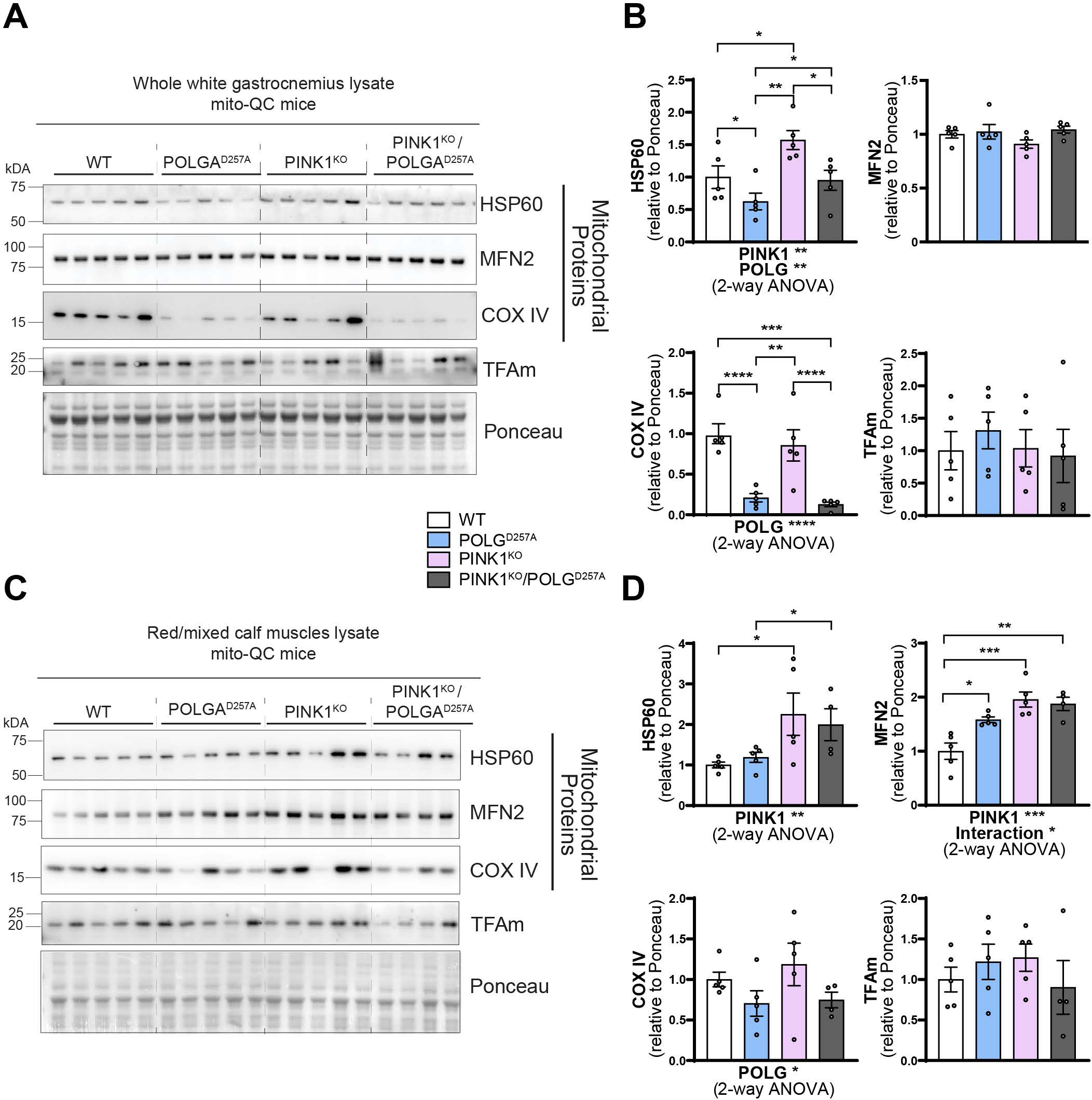
PINK1^KO^ increases mitochondrial content in skeletal muscles without affecting mitochondrial biogenesis. (A) Immunoblots of the indicated mitochondrial proteins and protein involved in mitochondrial biogenesis in the white gastrocnemius (WGC) of WT, PINK1 knock-out, mutator and double mutant (PINK1^KO^/POLG^D257A^) *mito*-QC mice. Quantitation (n=5 per group) of the (B) mitochondrial proteins and TFAm in the white gastrocnemius (WGC) displayed in (A). (C) Immunoblots of the indicated mitochondrial proteins and proteins involved in mitochondrial biogenesis in the red/mixed calf muscles (RCM, red/mixed gastrocnemius + plantaris). (D) Quantitation (n=4-5 per group) of the mitochondrial proteins and TFAm in the red/mixed calf muscles (RCM, red/mixed gastrocnemius + plantaris). Overall data is represented as mean +/-SEM. Statistical significance is displayed as *p<0.05, **p<0.01, ***p>0.001, and ****p<0.0001.

The increase in mitochondrial proteins in the PINK1 animals is indicative of either enhanced mitochondrial biogenesis or reduced mitophagy. For the former, we blotted for the mitochondrial transcription factor TFAm (Figure 2A & B). This protein is the master regulator of mitochondrial biogenesis as it controls mtDNA copy number and coordinates the expression of both nuclear and mitochondrial genomes [24,25]. TFAm levels were statistically unaffected, suggesting mitochondrial biogenesis is not responsible for the increase in mitochondrial markers, at least at this time point.

### Basal mitophagy is particularly impaired in PINK1 KO oxidative skeletal muscles

The above data suggests that mitophagy may be impaired in the PINK1 KO muscles and to determine if this was indeed the case, we used our *mito*-QC reporter to evaluate basal mitophagy levels in the WGC, RCM and in the soleus (Figure 3A & B, and Figure S3A). In all these muscle types, basal levels of mitophagy were detectable (red puncta only, detectable in all tissue). We observed that PINK1 KO moderately decreased mitophagy levels in WGC, but this effect was more pronounced in RCM and significant in the highly oxidative soleus. The presence of the POLG mutation only had a mild effect, with a subtle increase in mitophagy noted across all the samples compared to WT. However, there was a strong interaction effect between PINK1 KO and POLG^D257A^ mutations, as the double mutants displayed basal mitophagy levels similar to the WT group. The apparent rescue of the PINK1 KO mitophagy defect suggests that an independent mitophagy mechanism is upregulated upon the combined loss of PINK1 and POLG mutation. The BNIP3L/NIX pathway operates independently of PINK1 and Parkin [26] and has also been shown to compensate for loss of Parkin [27]. Consistent with a role for NIX in enhancing mitophagy, we found increased protein levels in the RCM of the double mutant (Figure 3C-F). However, further work is needed to confirm NIX-mediated mitophagy in this instance. Our results in skeletal muscles demonstrate that PINK1 ablation impairs basal mitophagy especially in the highly oxidative muscle, and that POLG mutation moderately increased in mitophagy independently of PINK1.

**Figure 3.**
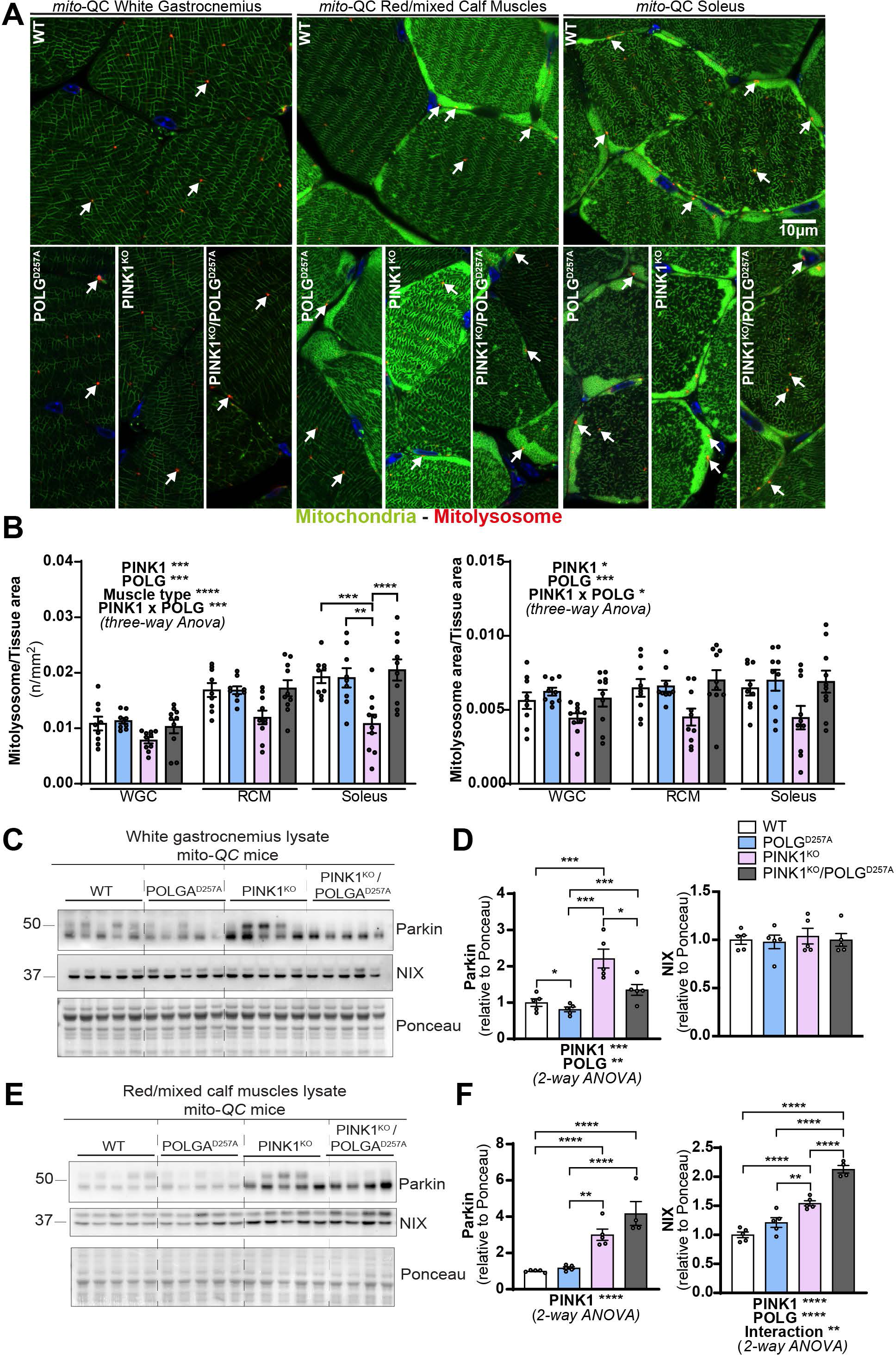
PINK1^KO^ decreases basal mitophagy in skeletal muscles, especially in oxidative fibres while POLG^D257A^ mildly increases it. (A) Representative micrographs of the white gastrocnemius (WGC), the red/mixed calf muscles (RCM, red/mixed gastrocnemius + plantaris), and soleus muscle of WT, PINK1 knock-out, mutator and double mutant (PINK1^KO^/POLG^D257A^) *mito*-QC mice. Scale bar: 10 µm. White arrows indicate examples of mitolysosomes. (B) Quantitation of basal mitophagy expressed as the number of mitolysosomes per tissue area, and the proportion of tissue occupied by mitolysosomes. (C) Immunoblots of the indicated proteins involved in different mitophagy pathways in the white gastrocnemius (WGC) of WT, PINK1 knock-out, mutator and double mutant (PINK1^KO^/POLG^D257A^) *mito*-QC mice. (D) Quantitation (n=5 per group) of Parkin, and BNIP3L/NIX in the white gastrocnemius (WGC) displayed in (C). (E) Immunoblots of the indicated proteins involved in different mitophagy pathways in the red/mixed calf muscles (RCM, red/mixed gastrocnemius + plantaris). Quantitation (n=4-5 per group) of Parkin, and BNIP3L/NIX in the red/mixed calf muscles (RCM, red/mixed gastrocnemius + plantaris) displayed in (E). Overall data is represented as mean +/-SEM. Statistical significance is displayed as *p<0.05, **p<0.01, ***p>0.001, and ****p<0.0001.

### Mitophagy in DA neurons

We had previously found little effect for PINK1 on basal mitophagy in other tissues [8], we wished to confirm that this was indeed the case in other tissues of the PINK1-POLG^D257A^ animals. Given that PD is a neurodegenerative disorder, characterized by a loss in SNpc DA neurons, and that DA neuron loss had been previously seen in older POLG^D257A^ mice where Parkin had been ablated [18], we focused on this area (Figure S3B-E). Firstly, we analyzed RNAseq data in the brain generated by the Genotype-Tissue Expression (GTEx) project. Despite PINK1 being relatively highly expressed in the substantia nigra, the rest of the pathway was not very highly expressed (Figure S3B) in comparison to our observations in skeletal muscles. At this age we noted no obvious reduction in tyrosine hydroxylase positive DA neuron numbers (Figure S3C & D). Unlike in skeletal muscles, and as previously observed, loss of PINK1 did not significantly affect basal mitophagy levels in these neurons, as quantified using the *mito*-QC reporter (Figure 3C & E). Additionally, there was a small but significant increase in mitophagy in mice harboring the POLG^D257A^ mutation, regardless of the PINK1 status. This suggests that in the mutator background, mitochondrial stress is increased, which is sufficient to mildly induce mitophagy. As with muscle, this is independent of PINK1.

## Discussion

Using genetics, we investigated the effects of PINK1 ablation in combination with the D257A mutation of DNA polymerase γ (POLG^D257A^) on mitophagy in mice of six months of age. We found an effect of PINK1^KO^ in oxidative skeletal muscle, suggesting this pathway is active within this tissue type under basal conditions. Interestingly, the presence of the POLG mutation only resulted in a moderate increase in mitophagy and any observed effects were independent of PINK1. Previously published work had shown increased DA neurodegeneration in mutator mice that lacked Parkin [18], suggesting that impaired PINK1/Parkin-dependent mitophagy could be responsible. While this conclusion may be at odds with our observations, we do note that animals here were much younger (6 months) compared to the previously published work (12 months), where the onset of the accelerated aging phenotype starts to manifest. It is therefore possible that the level of mitochondrial stress, caused by the POLG^D257A^ mutation, is below the “threshold” for PINK1 activation. It is also likely that tissues accumulate mutations at different rates with the POLG^D257A^ mutation and could possibly contribute to the heterogeneity of effects encountered in different organs. However, two recent publications have also failed to identify a neurodegenerative phenotype when the PINK1/Parkin pathway is lost in the background of the mutator mouse [28,29]. More work is obviously needed to determine the interaction of these pathways.

Regardless, we now show that PINK1 does have an impact on mitochondrial homeostasis and basal levels of mitophagy in skeletal muscles, in particular in the oxidative soleus. This is akin to observations in drosophila flight muscle [30]. Of note, the skeletal muscle is also the tissue in which PINK1 is proportionally the most highly expressed in humans, which could help explain our observations. We observed a gradient of alteration depending on muscle phenotype (glycolytic/oxidative) with only a stronger effect of PINK1 loss on mitophagy in the oxidative muscle. Our results thus highlight the importance of muscle phenotype in the study of the PINK1 pathway. Interestingly, physical activity has been shown to enhance symptoms of PD [31], hence maintaining skeletal muscle mitochondrial function seems to be a key factor in the aetiology of PD. It is possible that enhancing muscle and mitochondrial health would decrease peripheral inflammation, which is potentially responsible for a cascade of events leading to PD. However, as we do not know the significance of reduced mitophagy in PINK1 KO mouse muscle, or whether this is physiologically relevant to disease pathology, further work is needed. It is clear that multiple mitophagy pathways can operate within the same tissue, which could make determining the significance of one pathway in isolation more challenging.

As we can see evidence for impaired mitophagy upon loss of PINK1, this now gives us an opportunity to study the pathway in mammalian tissues, in the absence of overt stresses, for the first time. Not only will this allow an easier way to physiologically validate mechanistic findings from cell line studies, but also facilitate translational preclinical studies designed to target PINK1-dependent mitophagy therapeutically.

## Acknowledgements

We acknowledge Paul Appleton at the Dundee Imaging Facility, Dundee. We would also like to acknowledge Dr Lambert Montava Garriga, and the MRC Genotyping team for their expert technical assistance. The Zeiss LSM880 with Airyscan was supported by the ‘Wellcome Trust Multi-User Equipment Grant’ [208401/Z/17/Z]. This work was funded by grants to IGG from the Medical Research Council, UK (MC_UU_00018/2), and the Michael J. Fox Foundation (MJFF-003029).

## Material and methods

### RNA-seq data analysis

The data used for the analyses described in this manuscript were obtained from: Xena Browser data hubs [32] including 7433 samples from GTEx (Genotype-Tissue Expression) on 05/09/2023. We retained only tissue types that had a minimum of 20 samples available and applied quality control measures based on the distribution of gene expression for each sample. In the end, a total of 5,851 samples were chosen. Then, we normalized the gene expression data by doing quantile normalization. Codes can be found at: https://github.com/Keviosta/Singh_et_al. The Genotype-Tissue Expression (GTEx) Project was supported by the Common Fund of the Office of the Director of the National Institutes of Health, and by NCI, NHGRI, NHLBI, NIDA, NIMH, and NINDS.

### Animals

Experiments were performed on mice genetically altered in which the gene encoding for PTEN induced kinase 1 (PINK1) has been ablated, and/or carrying the D257A mutation of DNA polymerase subunit gamma (POLG^D257A^). The mitophagy (mito-QC) reporter mouse model used in this study was generated as previously described[14]. Experiments were performed on 42 adult mice (180 days old) of both genders (n=8–12 per group) all homozygous for the *mito*-QC reporter.

Animals were housed in sex-matched littermate groups of between two and five animals per cage in neutral temperature environment (21° ± 1°C), with a relative humidity of 55–65%, on a 12:12 hr photoperiod, and were provided food and water ad libitum. All animal studies were ethically reviewed and performed in agreement with the guidelines from Directive 2010/63/EU of the European Parliament on the protection of animals used for scientific purposes, and in accordance with the Animals (Scientific Procedures) Act 1986. All animal studies and breeding were approved by the University of Dundee ethical review committee, and further subjected to approved study plans by the Named Veterinary Surgeon and Compliance Officer and performed under a UK Home Office project license in agreement with the Animal Scientific Procedures Act (ASPA, 1986).

### Sample collection

Mice were terminally anesthetised with an intraperitoneal injection of a pentobarbital sodium solution (400 mg/kg, Euthatal, Merial) prepared in PBS (Gibco, 14190–094), then trans-cardially perfused with DPBS (Gibco, 14190–094) to remove blood. Tissues were rapidly harvested and either snap frozen in liquid nitrogen and stored at −80°C for later biochemical analyses or processed by overnight immersion in freshly prepared fixative: 3.7% Paraformaldehyde (Sigma-Aldrich, 441244) 200 mM HEPES, pH=7.00. The next day, fixed tissues were washed three times in DPBS, and immersed in a sucrose 30% (w/v) solution containing 0.04% sodium azide until they sank at the bottom of the tube. Samples were stored at 4°C in the aforementioned sucrose solution until further processing.

### Tissue sectioning

Tissues were embedded in an optimal cutting temperature compound matrix (O.C.T., Scigen, 4586), frozen and sectioned with a Leica CM1860UV cryostat using carbon-steel microtome blades (Feather, C35). Twelve microns sections were placed on slides (Leica Surgipath X-tra Adhesive, 3800202), air dried, and then kept at −80°C in a slide box containing a silica gel packet (Sigma-Aldrich, S8394) until further processing. Sections were thawed at room temperature and washed three times for 5 minutes in DPBS (Gibco, 14190–094). Sections were then counterstained for 5 minutes with Hoechst 33342 (1μg/mL, Thermo Scientific, 62249). Slides were mounted using Vectashield Antifade Mounting Medium (Vector Laboratories, H-1000) and high-precision cover glasses (No. 1.5H, Marienfeld, 0107222) and sealed with transparent nail polish.

### Brain free-floating sectioning and immunolabeling

50-µm thick brain free-floating sections were obtained by axial sectioning using a sliding microtome (Leica, SM2010R) equipped with a dry-ice tray (Leica 14050842641). Sections were collected using a round paintbrush, placed in DPBS (Gibco, 14190–094), and stored at 4°C until further use. Free-floating sections were permeabilised using DPBS (Gibco, 14190– 094) containing 0.3% Triton X-100 (Sigma Aldrich, T8787) three times for 5 min. Sections were then blocked for 1 hr in blocking solution (DPBS containing 10% goat serum (Sigma Aldrich, G9023), and 0.3% Triton X-100). Overnight primary antibody incubation was performed on a rocker in blocking solution containing the Anti-Tyrosine Hydroxylase (1/1000, Millipore, AB152) antibody. The next day, sections were washed two times for 8 min in DPBS containing 0.3% Triton X100 and then incubated for 1 hr in blocking solution containing the secondary antibody (1/200, Invitrogen P10994 Goat anti-Rabbit IgG (H+L) Cross-Adsorbed Secondary Antibody, Pacific Blue). Sections were then washed two times for 8 min in DPBS and mounted on slides (Leica Surgipath X-tra Adhesive, 3800202) using Vectashield Antifade Mounting Medium (Vector Laboratories, H-1000) and sealed with nail polish.

### Confocal microscopy

Confocal micrographs were obtained by uniform random sampling using a Zeiss LSM880 with Airyscan laser scanning confocal microscope (Plan-Apochromat 63x/1.4 Oil DIC M27; Carl Zeiss Immersol immersion oil 518F, 444960-0000-000) using the optimal resolution for acquisition (Nyquist sampling), averaging 4, bit depth 16 bits, and with a pixel dwell of 1.10 µs. 10–15 images were acquired per sample, depending on the tissue, by an experimenter blind to all conditions. High-resolution, representative images were obtained using the Super Resolution mode of the Zeiss LSM880 with Airyscan. Representative images were altered uniformly within the same figure by increasing intensity levels and/or by applying a median filter (half size = 2) to enhance visualisation, using the Icy software [33].

### Quantitation of mitophagy

Images were processed with Volocity Software (version 6.3, Perkin-Elmer) as previously described [16]. Briefly, images were first filtered using a fine filter to suppress noise. Tissue was detected by thresholding the Green channel. For the immunolabelings in the brain, dopaminergic neurons of the substantia nigra pard compacta were detected by thresholding the Pacific Blue labelled channel. A ratio image of the Red/Green channels was then created for each image. Mitolysosomes were then detected by thresholding this ratio channel as objects with a high Red/Green ratio value within the tissue/cell population of interest. The same ratio channel threshold was used per organ/set of experiments. To avoid the detection of unspecific high ratio pixels in the areas of low reporter expression, a second threshold of the red channel was applied to these high ratio pixels. This double thresholding method provides the most accurate detection of mitolysosomes as structures with a high Red/Green ratio value and a high Red intensity value.

### Preparation of protein lysates and western blotting

Samples were processed from 5 randomly selected animals per group to minimise the effect of interindividual variability. For further consistency, the same animals and loading order were used in all blots from different tissues. Due to a sample issue in the RCM, the PINK1^KO^/POLG^D257A^ group only contains 4 samples.

Frozen tissue was disrupted with a liquid-nitrogen cooled Cellcrusher (Cellcrusher, Cork, Ireland) tissue pulveriser. Approximately 20–30 mg of pulverised tissue were then lysed on ice for 30 min with (10 µL/mg tissue) of RIPA buffer [50 mM Tris–HCl pH 8, 150 mM NaCl, 1 mM EDTA, 1% NP-40, 1% Na-deoxycholate, 0.1% SDS, and cOmplete protease inhibitor cocktail (Roche, Basel, Switzerland)], phosphatase inhibitor cocktail (1.15 mM sodium molybdate, 4 mM sodium tartrate dihydrate, 10 mM β-glycerophosphoric acid disodium salt pentahydrate, 1 mM sodium fluoride, and 1 mM activated sodium orthovanadate), and 10 mM DTT. All inhibitors and DTT were added immediately prior to use. Crude lysates were vortexed and centrifuged for 10 min at 4°C at 20,817xg. The supernatant was collected, and the protein concentration determined using the Pierce BCA protein assay kit (ThermoFisher Scientific, Waltham, MA, USA). For each sample, 8 µg of protein was separated on a NuPAGE 4–12% Bis-Tris gel (Life technologies, Carlsbad, CA, USA). Proteins were electroblotted to 0.45 µm PVDF membranes (Imobilon-P, Merck Millipore, IPVH00010; or Amersham Hybond, GE Healthcare Life Science, 10600023), and immunodetected using primary antibodies directed against HSP60 rabbit polyclonal (D307) (1/1000, Cell Signalling Technology, #4870S), MFN2 rabbit monoclonal (1/1000, Abcam, ab124773), COX IV rabbit monoclonal (1/1000, Cell Signalling Technology, #4850S), TFAm mouse monoclonal [18G102B2E11] (1/1000, abcam, ab131607), Parkin mouse monoclonal (1/1000, Santa Cruz Biotechnology, sc-32282), BNIP3L/NIX (D4R4B) rabbit monoclonal antibody (1/2000, Cell Signalling Technology, #12396S). Protein revelation was performed using HRP-conjugated secondary antibodies, using ECL substrates (Bio-Rad, Clarity, 1705061; or Clarity Max, 1705062), and a ChemiDoc MP (Bio-Rad) imaging system. Image analysis was performed with FIJI [34]. As we were not able to find an appropriate housekeeping gene that did not vary with our experimental conditions in the skeletal muscles, whole lane Ponceau was used for normalization.

### PINK1 immunoprecipitation

For immunoprecipitation of endogenous PINK1, 500 μg of whole-brain lysate was incubated overnight at 4°C with 10 μg of PINK1 antibody (S774C; MRC PPU Reagents and Services) coupled to Protein A/G beads (10 μl of beads per reaction; Amintra) as previously reported [8]. The immunoprecipitants were washed three times with lysis buffer containing 150 mM NaCl and eluted by resuspending in 10 μl of 2× LDS sample buffer and incubating at 37°C for 15 min under constant shaking (2000 rpm) followed by the addition of 2.5% (by volume) 2-mercaptoethanol.

### Statistics

Data are represented as means ± SEM. Number of subjects are indicated in the respective figure legends. Statistical analyses were performed using a two-way analysis of variance (ANOVA) or three-way ANOVA followed by a Tukey multiple comparison test using GraphPad Prism for Windows 64-bit version 9.5.1 (733) (GraphPad Software, San Diego, California USA). Statistical significance is displayed as * p< 0.05: ** p < 0.01, *** p<0.001, and **** p<0.0001.

---

**Figure S1.**
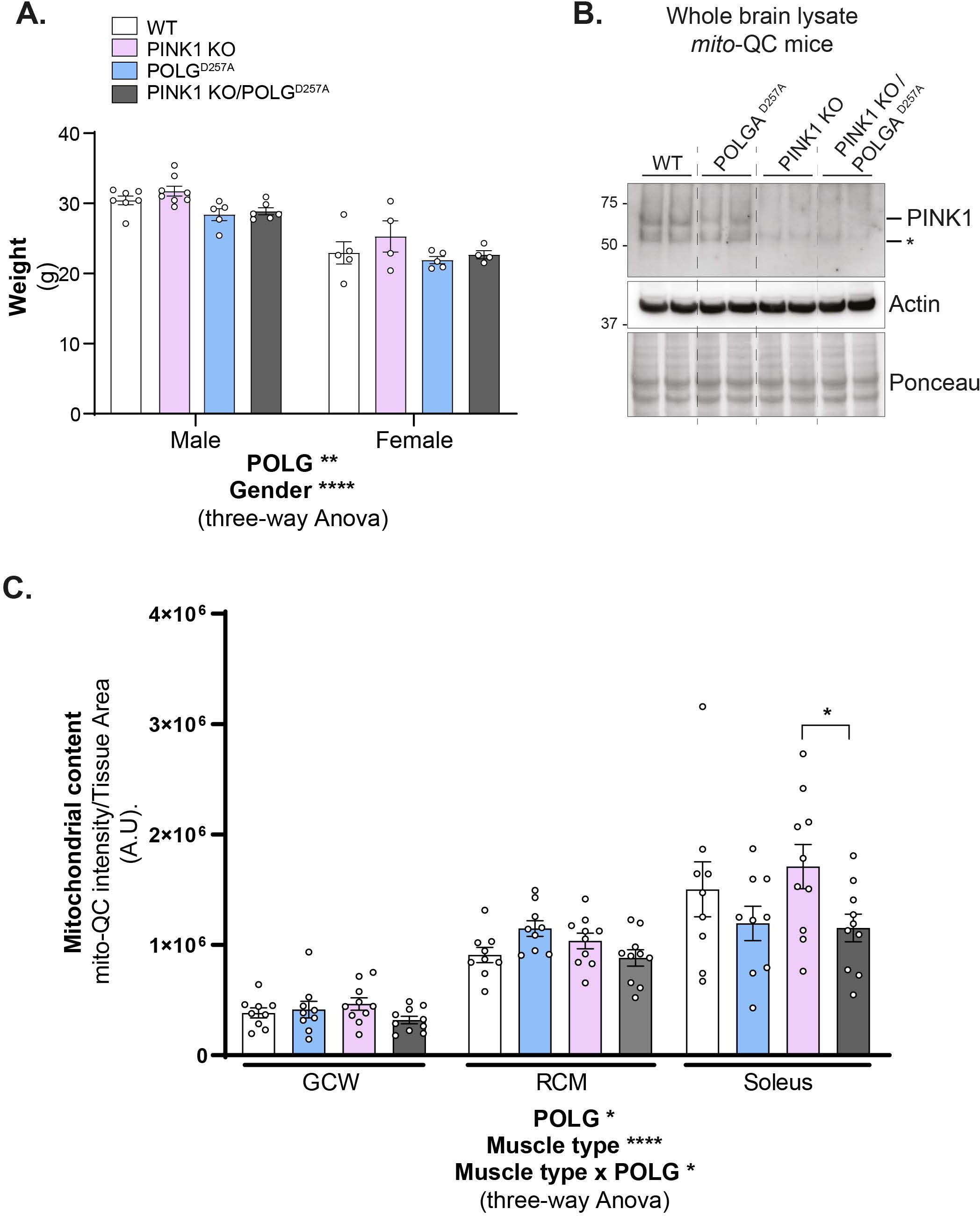
(A) Body weight of 180 days old mice of WT, PINK1 knock-out, mutator and double mutant (PINK1^KO^/POLG^D257A^) *mito*-QC mice (n=10-12 per group). Males and females were separated to better visualize the data. (B) Immunoblot of PINK1 immunoprecipitation from whole brain lysates of the same 4 groups (n=2 per group). Asterisk indicates an unspecific band. (C) Relative mitochondrial content in each muscle phenotype determined using the *mito*-QC GFP expression. (n=9-10). Overall data is represented as mean +/-SEM. Statistical significance is displayed as *p<0.05, **p<0.01, and ****p<0.0001.

**Figure S3.**
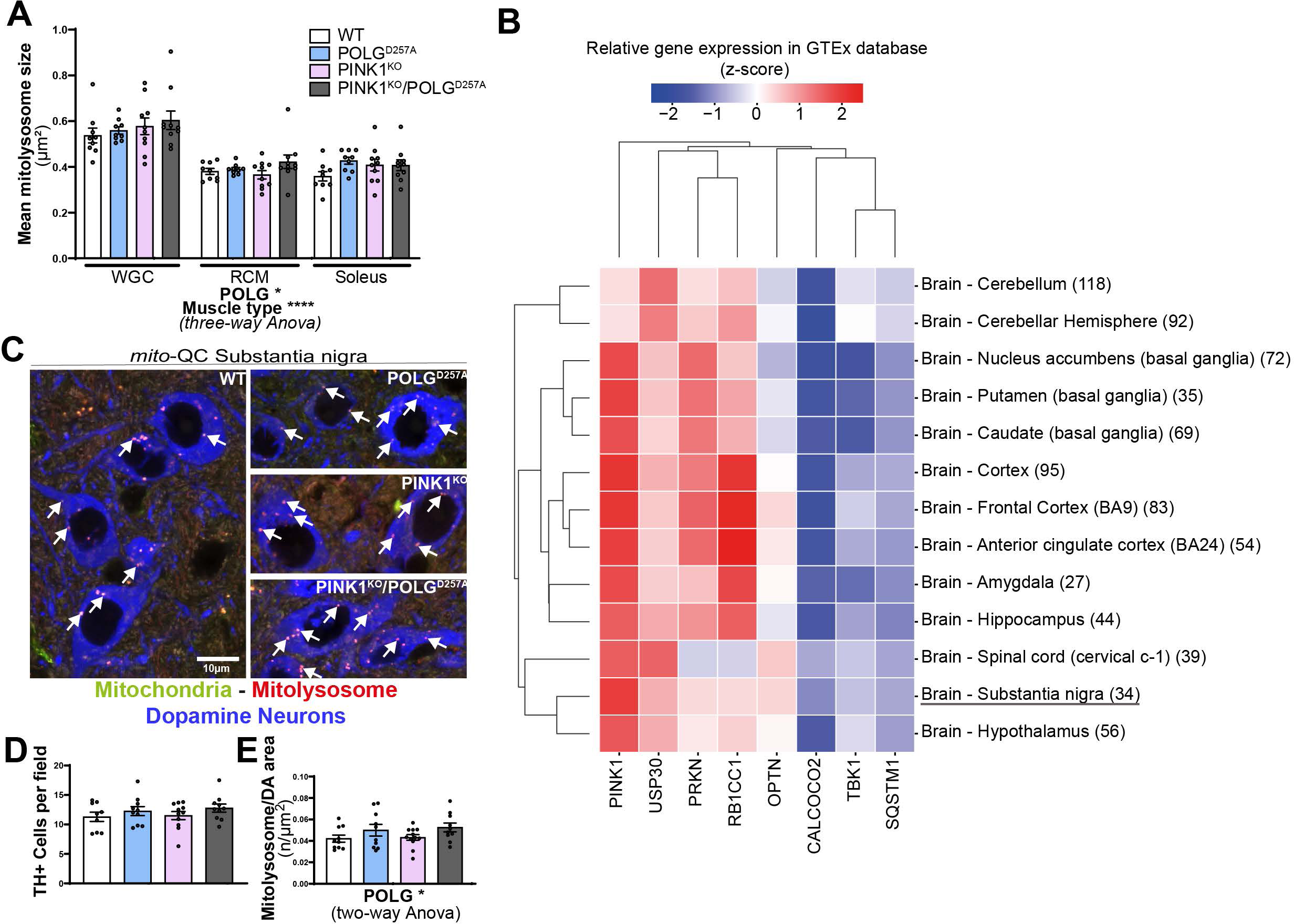
(A) Quantitation of basal mitophagy expressed by average size of a mitolysosome in the different areas of the skeletal muscles. (B) Gene-expression RNA-seq data analysis of the PINK1/Parkin pathway in different human brain regions, obtained from GTEx (Genotype-Tissue Expression). (C) Representative micrographs of tyrosine hydroxylase (TH) immunolabelled dopaminergic neurons of the substantia nigra pars compacta in WT, PINK1 knock-out, mutator and double mutant (PINK1^KO^/POLG^D257A^) *mito*-QC mice. Scale bar: 10 µm. White arrows indicate examples of mitolysosomes. (D) Quantitation of the number of TH positive cells per field of view using a 63x objective. (E) Quantitation of basal mitophagy in the DA neurons of the SNpc expressed as the number of mitolysosomes per TH positive area, (n=10-12). (Overall data is represented as mean +/-SEM. Statistical significance is displayed as *p<0.05, and ***p<0.0001.

